# Landscape permeability and individual variation in a dispersal-linked gene jointly determine genetic structure in the Glanville fritillary butterfly

**DOI:** 10.1101/409003

**Authors:** Michelle F. DiLeo, Arild Husby, Marjo Saastamoinen

## Abstract

There is now clear evidence that species across a broad range of taxa harbour extensive heritable variation in dispersal. While studies suggest that this variation can facilitate demographic outcomes such as range expansion and invasions, few have considered the consequences of intraspecific variation in dispersal for the maintenance and distribution of genetic variation across fragmented landscapes. Here we examine how landscape characteristics and individual variation in dispersal combine to predict genetic structure using genomic and spatial data from the Glanville fritillary butterfly. We used linear and latent factor mixed models to identify the landscape features that best predict spatial sorting of alleles in the dispersal-related gene *phosphoglucose isomerase* (*Pgi*). We next used structural equation modeling to test if variation in *Pgi* mediated gene flow as measured by F_st_ at putatively neutral loci. In a year when the population was expanding following a large decline, individuals with a genotype associated with greater dispersal ability were found at significantly higher frequencies in populations isolated by water and forest, and these populations showed lower levels of genetic differentiation at neutral loci. These relationships disappeared in the next year when metapopulation density was high, suggesting that the effects of individual variation are context dependent. Together our results highlight that 1) more complex aspects of landscape structure beyond just the configuration of habitat can be important for maintaining spatial variation in dispersal traits, and 2) that individual variation in dispersal plays a key role in maintaining genetic variation across fragmented landscapes.

**Impact summary:** Understanding how fragmentation affects dispersal and gene flow across human-modified landscapes has long been a goal in evolutionary biology. It is typically assumed that individuals of the same species respond to the landscape in the same way, however growing evidence suggests that individuals can vary considerably in their dispersal traits. While the effects of this individual dispersal variation on range expansions and invasions have been well-characterized, knowledge of how it might mediate genetic responses to landscape fragmentation are almost entirely lacking. Here we demonstrate that individual variation in dispersal is key to the maintenance of genetic variation during a population expansion following a large decline in a butterfly metapopulation. We further show that spatial variation in dispersal is not maintained by the configuration of habitat patches alone, but by a more complex genotype-environment interaction involving the landscape matrix (i.e. landscape features found between habitat patches). This challenges the simplified landscape representations typically used in studies of dispersal evolution that ignore heterogeneity in the landscape matrix. More broadly, our results highlight the interplay of adaptive and neutral processes across fragmented landscapes, suggesting that an understanding of species vulnerability to landscape fragmentation requires consideration of both.

## Introduction

Dispersal is key to the maintenance of genetic variation and adaptive potential in fragmented landscapes. Differences in species ability to maintain genetic diversity in fragmented landscapes can, in part, be explained by interspecific differences in dispersal capacity (Steele *et al.* 2009; Peterman *et al.* 2015). However, there is now clear evidence that many species across a broad range of taxa harbour extensive heritable variation in dispersal (Saastamoinen *et al.* 2018), which can facilitate demographic outcomes such as range expansion (e.g. Duckworth and Badyaev 2007; Ochocki and Miller 2017) and invasions (e.g. Phillips *et al.* 2006; Elliott and Cornell 2012; Cote *et al.* 2017). However, few studies have considered the consequences of intraspecific variation in dispersal for genetic outcomes such as the maintenance and distribution of genetic variation across fragmented landscapes (Cheptou *et al.* 2017). This is an important gap given that the genetic makeup of populations can drive the trajectories of both ecological and evolutionary processes (Rius and Darling 2014; Szucs *et al.* 2017; Wagner *et al.* 2017).

This gap comes partly from a lack of integration of intraspecific variation into fields like landscape and spatial genetics (Edelaar and Bolnick 2012; Pflueger and Balkenhol 2014). Studies in these fields emphasize that gene flow across fragmented landscapes is strongly constrained by population configuration, habitat quality, and matrix heterogeneity (i.e. the landscape features intervening populations), but typically assume that dispersers respond to the landscape in the same way (Manel *et al.* 2003; Holderegger and Wagner 2008). However, fragmentation itself can exert strong selective pressures on dispersers, in some cases leading to the maintenance of multiple dispersal strategies across the same landscape (Cheptou *et al.* 2017; Cote *et al.* 2017; Legrand *et al.* 2017). This variation might change expectations of spatial genetic structure and mask landscape genetic relationships, yet empirical tests of this are lacking (but see McDevitt *et al.* 2013). For example, using simulations, Palmer *et al.* (2014) showed that distance-based connectivity metrics underestimated the number of migrants arriving into isolated populations when individuals were allowed to vary in their dispersal ability, with the effect most severe when dispersal was rare. This spatial sorting of good dispersers into more isolated populations is expected to also impact the distribution of genetic variation, for example by increasing genetic neighbourhoods beyond expectations drawn from mean dispersal distances (DiLeo *et al.* 2014). Because dispersal traits often co-evolve with other aspects of morphology, physiology, and behaviour (Clobert *et al.* 2009; Cote *et al.* 2017), individuals might also interact with, and respond to, the landscape matrix in different ways (Merckx and Van Dyck 2007; Delgado *et al.* 2010). This could mean that the effects of landscape on gene flow might be missed if intraspecific variation is ignored.

Intraspecific variation thus has the potential to play an important, but so far unexplored role in structuring species genetic response to landscape fragmentation. The maintenance of several dispersal strategies across a single landscape might allow wide-spread gene flow to be maintained under a broad set of ecological conditions. We test this in a model species, the Glanville fritillary butterfly (*Melitaea cinxia*) in the Åland Islands, Finland. Importantly, individuals vary in their dispersal ability: butterflies heterozygous or homozygous for the “c” allele in a SNP associated with the gene *phosphoglucose isomerase* (*Pgi*) have higher flight metabolic rate, which translates to substantially increased dispersal propensity and dispersal distance in the field, especially at cooler temperatures (reviewed in Niitepõld and Saastamoinen 2017). Evidence for this comes from laboratory experiments linking flight metabolic rate to *Pgi* genotype (Niitepõld 2010), and from a study linking dispersal distances of butterflies tracked in the field to *Pgi* genotype (Niitepõld *et al.* 2009). The butterfly persists in a highly dynamic metapopulation with frequent colonizations and extinctions, and we focus on a two-year period representing extremes of fluctuations experience by the butterfly: the year 2011 where populations have extremely low connectivity because of a large decline in the previous year, and 2012 where populations have high connectivity following the recovery of populations in the year prior (Ojanen *et al.* 2013). Recent work suggests that *Pgi* plays a central role in metapopulation persistence by maintaining high recolonization rates despite drastic population fluctuations (Hanski *et al.* 2017). However, we do not yet know to what extent *Pgi* also contributes to the maintenance of neutral genetic variation, which is important given that genetic diversity can exert effects on persistence independently of demographic rescue (e.g. Szucs *et al.* 2017). Specifically, we ask: 1) what landscape factors drive spatial sorting of genotypes that vary in their dispersal ability? And, 2) does this dispersal variation mediate the genetic response to landscape structure and population fluctuations? While previous work has found that more isolated patches have higher frequencies of good dispersers (Haag *et al.* 2005, Hanski and Saccheri 2006; Zheng *et al.* 2009), we further predict that dispersers will respond differentially to heterogeneity in the landscape matrix. We also predict that the good dispersers will facilitate genetic admixture; population with higher frequencies of the *Pgi-c* allele should be less genetically differentiated than populations with low frequencies. Finally, we predict that the effects of *Pgi* will be context dependent. Modeling predicts that the dispersive genotype will have the largest advantage when there are many open patches to colonize (Zheng *et al.* 2009), and thus we expect the association between landscape structure, *Pgi*, and genetic structure to be highest in 2011.

## Methods

### Study species and sampling

In Finland, the Glanville fritillary butterfly is only found in the Åland Islands where it persists in a metapopulation encompassing over 4000 meadows (hereinafter ‘patches’) that contain one or both of its host plants, *Plantago lanceolata* and *Veronica spicata*. In late summer, females lay clutches of 150-200 eggs, which develop into larvae that live gregariously until the last larval instar in the following spring (Boggs and Nieminen 2004). Before winter diapause, larvae spin nests at the base of host plants, and every fall since 1993 the number of nests have been counted in all patches in Åland allowing the quantification of long-term population dynamics (see Ojanen *et al.* 2013 for survey methods). In any year, only about 20% of patches on average are occupied, with frequent local recolonizations and extinctions, and large variation in the number of total larval nests (Fig. 1). Our study focused on the period 2010-2012, which is characterized by a large population decline in 2010 due to poor weather conditions, followed by a population expansion. The major expansion occurred in 2011 where a record number of new colonizations was documented (Fig. 1b). In 2012, there were fewer colonization of previously empty patches but a large increase in population density, with a record number of larval nests (Fig 1a).

**Figure 1.**
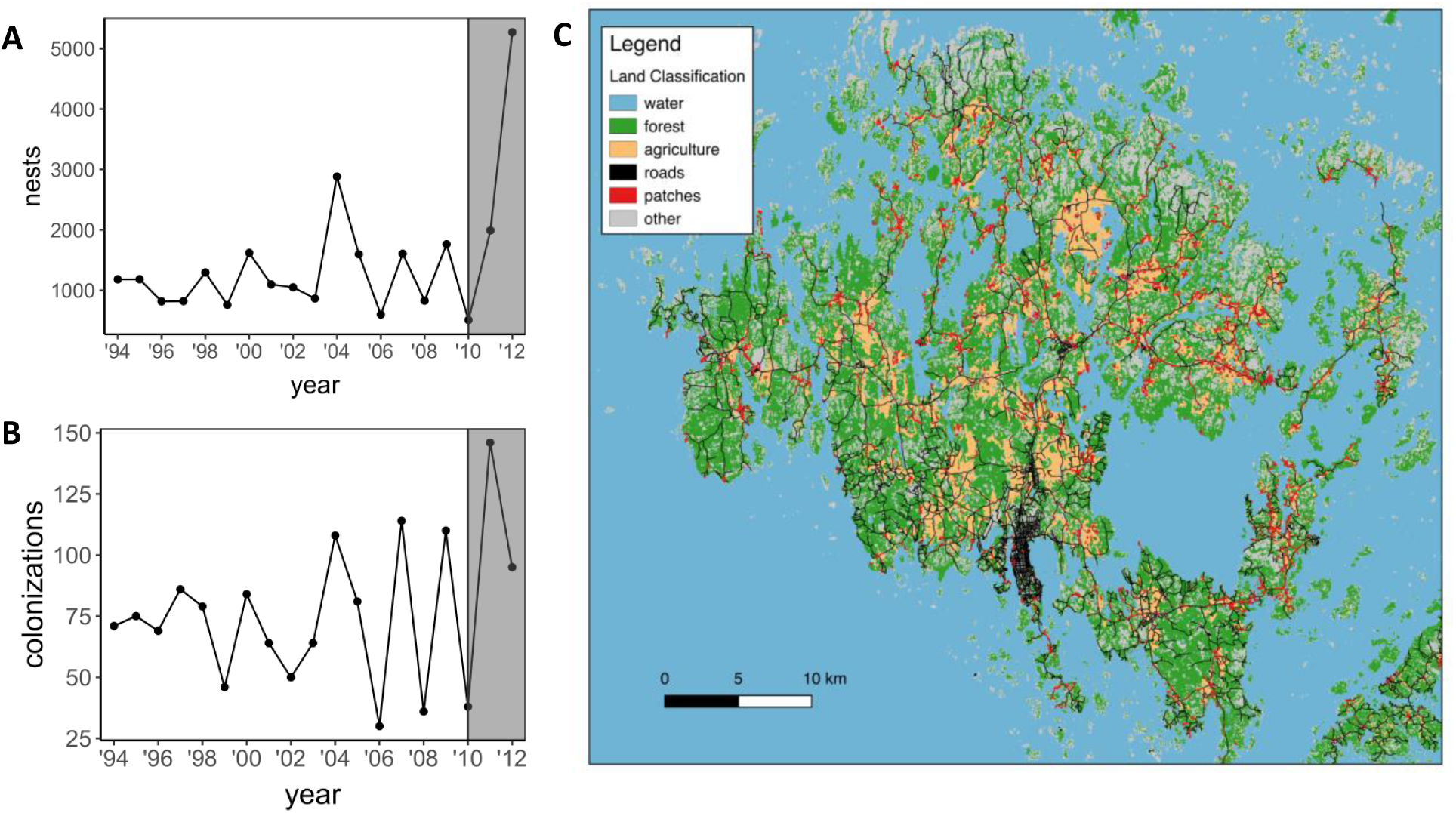
Yearly variation in the number of nests (a) and colonizations (b) in the Åland Islands (c). The period of study from 2010-2012 is highlighted in a-b, and c shows the major land cover types across the study region. Data for population trends come from annual surveys described in detail in Ojanen *et al.* 2013.

In 2011 and 2012, three larvae per nest were collected from patches across Åland for genetic analysis. Larval tissue was homogenized and DNA was extracted as described in Fountain *et al.* (2018). Genotyping was performed on the KASP platform. In total 40 putatively neutral markers from noncoding regions of the genome were selected, and an additional five putatively functional markers were selected from candidate genes related to flight or dispersal traits including *Pgi* (Orsini *et al.* 2009; Kvist *et al.* 2013; Kvist *et al.* 2015; Wong *et al.* 2016; Duplouy *et al.* 2017; Appendix B). The neutral loci were obtained from SOLiD matepair-1 genome sequences (Ahola *et al.* 2014) using an in-house SNP calling method (Rastas *et al.* 2013). The neutral SNPs have minor allele frequencies >0.2, and span all 31 chromosomes from non-coding regions. Further details of SNP calling, validation, and quality control are described in Fountain *et al.* (2016). Previous work demonstrated that this neutral SNP panel was sufficient to capture both large (Nair *et al*. 2016) and small-scale (Fountain *et al.* 2018) patterns of genetic population structure. In a study conducted in a smaller region in Åland, Fountain *et al.* (2018) showed that the neutral 40-SNP panel and a larger panel of 272 SNPs resolved the same patterns of spatial genetic structure in each of the six years tested. We are thus confident that the panel used here will represent the genetic structure of the metapopulation well.

After excluding any SNP or individual with a call rate <0.95 percent, 34 neutral makers and all of the functional markers remained, and total sample sizes were 3365 larvae representing 1504 families in 250 patches in 2011, and 8229 larvae representing 2999 families in 322 patches in 2012. We only had a few samples from the extreme south of our study region in Lemland. As these samples were clear outliers with low connectivity and frequencies of *Pgi*-c (Figs. S1-S2), we excluded them from downstream analysis. We further excluded 98 patches in 2011 and 129 in 2012 with only a single larval nest where the effects of genetic drift are expected to be especially strong, giving a sample size of 152 patches (74 old, 78 new) in 2011 and 193 patches (138 old, 55 new) in 2012.

### Development of landscape connectivity hypotheses

To test the effects of landscape on neutral and functional genetic variation, we developed a set of landscape resistance surfaces reflecting the permeability of the intervening landscape matrix to dispersing butterflies. The landscape was classified from 20 m resolution CORINE 2012 Land Cover Inventory raster data from the Finnish Environmental Institute (http://www.syke.fi/en-US/Open_information/Spatial_datasets). The original raster layer included 48 land cover categories, 42 of which were found in our study region. We additionally overlaid polygons representing patches, and added a category representing edges between agriculture and forested areas as previous work found that *M. cinxia* tend to move along edges (Ovaskainen *et al.* 2008). We simplified the final land classification by combining structurally similar landscape features, resulting in 11 distinct categories representing major land use in the study region: discontinuous urban, continuous urban, open water, closed forest, transitional woodland scrub, agriculture with no major natural elements, agricultural edges, pasture, bare rock, roads, and patches.

We assigned each of these features a value representing its resistance to a dispersing *M. cinxia* individual. We assigned each feature either a value of one (not resistant to movement) or 10 (restricts movement), generating a total of 20 surfaces with different combinations of resistant and non-resistant features (Table S1). Patches were always given a value of one since patches represent suitable habitat for the species, and continuous urban areas were always given a value of 10. For each of these surfaces, we calculated pairwise resistance distances between *M. cinxia* patches using the program `CIRCUITSCAPE` (McRae 2006). This program uses circuit theory to calculate effective distances among patches by taking into account the relative permeability of the intervening landscape.

For each of the landscape hypotheses, we calculated an index of connectivity for each occupied patch using the incidence function model (Hanski 1994):

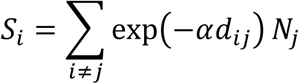

where *d*_*ij*_ is either the geographic distance (in km) or resistance distance (from `CIRCUITSCAPE`) between focal patch *i* occupied in year *t* and source patch *j* occupied in the previous year (*t*-1), and *N*_*j*_ is the number of *M. cinxia* nests found in source patch *j* in year *t*-1. The constant α scales the dispersal kernel and should be equal to 1/mean dispersal distance of the species, which has been estimated to be 1 km (Fountain *et al.* 2018). Because resistance distances are unit-less, mean dispersal distance for each landscape resistance hypothesis was chosen as the resistance value that roughly translated to 1 km in Euclidean distance units, which was then used to calculate α (Table S1). We did this by taking the predicted value of the resistance distance at a Euclidean distance value of 1 km using a simple linear model, after first linearizing relationships using log transformations. Many of the connectivity variables calculated from alternative landscape resistance surfaces were highly correlated, because some landscape features were only present in small proportions in our study region and thus did not have large effects on the calculations of resistance distances. For our main analyses we settled on five uncorrelated variables of landscape connectivity: Si_metapop_, which is scaled by Euclidean distance among patches but assumes that the intervening landscape does not restrict dispersal, Si_water_, Si_forest_, and Si_agriculture_, which assume that water, forest, and agriculture restrict dispersal, respectively, and Si_roads_, which assumes that roads facilitate dispersal. The five connectivity variables had pearson correlations below 0.6 and variance inflation factors in linear models under two (Table S2).

### Associations between landscape connectivity and Pgi

Our first objective was to test which landscape connectivity hypothesis best predicted the distribution of the *Pgi*-c allele. Based on previous work, we expected an interactive effect of patch age; good dispersers are expected to be at the highest frequency in newly colonized (hereinafter “new”), isolated populations and at the lowest in old, isolated populations because dispersive individuals also have high emigration rates (Zheng *et al.* 2009). We conducted model selection on mixed effect models including all connectivity variables and their interaction with patch age (i.e. new or old population) as fixed effects, and genetic cluster membership as a random effect. Frequency of the *Pgi-c* allele in each patch was used as the response variable. Genetic cluster membership represents groups of individuals or populations that share a common demographic history (i.e. share common historical population dynamics and/or ancestry), and was determined for each year separately using BAPS6 (see Appendix A; Fig. S1; Corander *et al.* 2008; Cheng *et al.* 2013). Mixed models were run for each year separately and we applied log transformations to *Si*_*metapop*_ and to *Si*_*roads*_, and exponential transformations to *Si*_*water*_ and *Si*_*agriculture*_ to linearize relationships. Each variable was scaled and mean centred. Models were implemented in the lme4 library (Bates *et al.* 2014) and validated graphically by plotting residuals against fitted values and normality assumptions were checked with QQ plots. For model selection, we retained a candidate set of models with high support for further analysis and interpretation. A model was included in the candidate set based on two conditions: 1) the model was within ΔAICc < 2 of the top model, and 2) the model was not simply an embellishment of a higher ranked model (i.e. did not contain uninformative parameters; Arnold 2010). All predictors appearing in the resulting candidate model set were considered as potentially important in their effects on *Pgi* and were subject to downstream analysis (see below). We used this approach since alternatives such as model averaging and summing akaike weights can lead to flawed interpretation of effects when variables are even weakly collinear (Galipaud *et al.* 2014; Cade 2015).

The residuals of the top model for both 2011 and 2012 showed evidence of spatial autocorrelation. To test that this did not bias our results, we compared models with and without spatial random effects implemented in r-inla (Lindgren & Rue 2015; Appendix A). In one case we found the sign of a weak main effect changed in the spatial compared to the non-spatial model (from 0.006 to - 0.003; Table S3). However, the major effect of this variable manifested as an interaction with a strong negative association in new populations, and the direction and strength of this interaction did not change. We found very little difference in all other estimates between the spatial and non-spatial models, and thus did not pursue spatial models further (Table S3).

As a second step, we used gene-environment association analysis (Rellstab *et al.* 2015) using latent factor mixed models (lfmms; Frichot *et al.* 2013) implemented in the LEA library (Frichot and François 2015) to confirm associations identified as significant in the linear mixed effect models. While the linear mixed model approach controlled for potential effects of neutral genetic structure by including genetic cluster membership (calculated from the 34 neutral loci) as a random effect, the latent factor mixed model approach explicitly considers all loci (*Pgi* and neutral loci) simultaneously. This approach applies a stronger control of background genetic structure, particularly when structure is complex and hierarchical (de Villemereuil *et al.* 2014), and also allowed us to confirm that we only see associations between connectivity and allele frequencies for *Pgi* and not for other loci. A drawback is that it cannot incorporate additive or interactive effects, and thus we used it only to confirm results in the partitions of the data that were found to be significant in linear mixed effect models (e.g. new populations). In addition to testing the individual Si variables identified as important in the mixed models, we explored potential additive effects by testing for association between loci and Si variables that incorporated the resistance of two or more of the landscape features. For example, the additive effects of Si_water_ and Si_forest_ were included in the model as the predictor Si_water+forest_ (Table S1).

Appendix A describes how the latent factors were specified starting from knowledge of the number of *k* genetic clusters identified from `BAPS6`. We ran the lfmms 10 times for each connectivity predictor, for 20,000 iterations following 10,000 iterations of burn-in. Loci included in each analysis were *Pgi*, 34 neutral markers, and four additional markers previously identified as being outliers in fragmented versus continuous landscapes (Fountain *et al.* 2016). We calculated median z-scores across the 10 repetitions. The resulting *p-*values were calibrated to correct for type I errors by applying an inflation factor (Francois *et al.* 2016). We corrected for multiple testing by applying the Benjamini-Hochberg algorithm (Benjamini and Hochberg 1995) with a false discovery rate of 10%.

### Direct and indirect effects of landscape structure on genetic structure

Our second objective was to test if intraspecific variation at the *Pgi* locus mediates the effect of landscape on the distribution of neutral genetic variation. We hypothesized that landscape structure will influence neutral genetic structure either directly (i.e. by limiting dispersal of all individuals in the same way), or indirectly through its effect on the spatial distribution of individuals with contrasting *Pgi* genotypes. We first tested for direct effects for each year separately, and then tested for indirect effects in partitions of the data where we found strong associations between connectivity and *Pgi* in latent factor mixed models.

Direct effects were tested using linear mixed effect models with neutral genetic differentiation as a response variable. A patch-specific measure of genetic differentiation was calculated by taking the average pairwise Weir and Cockerham unbiased F_st_ (Weir and Cockerham 1984) for each patch following Pflueger and Balkenhol (2014). Prior to calculation of F_st_, the data were subset to include only a single individual per family, and a maximum of 50 families per patch since there was large variation in sample sizes. Although Weir and Cockerham’s unbiased F_st_ accounts for unequal population sizes, calculations are biased at very low sample sizes (Willing *et al.* 2012), and so we limited analyses to patches with more than two families, excluding 43 patches in 2011 and 37 patches in 2012. Models were run for each year separately with the five connectivity metrics and population age as fixed predictors, and genetic cluster membership as a random factor. We did not find evidence for an interactive effect of population age and so only included it as a main effect. Predictor variables were transformed, centred, and scaled, and model selection was implemented as described above (see ‘Association between landscape connectivity and *Pgi*’).

We next tested for indirect effects of landscape structure on genetic differentiation in the partitions of data where we found association between connectivity and *Pgi* (see above) using structural equation modeling (SEM) implemented in the lavaan library (Rosseel 2012). Originally developed by Sewall Wright (1934), SEM allows the evaluation of complex *apriori-*defined relationships among variables that potentially involve direct and indirect effects. It is thus an ideal framework for testing the relative effects of connectivity versus *Pgi* on genetic structure, while controlling for directed or residual relationships between them. We modelled two pathways: (1) connectivity directly affects genetic differentiation, and (2) connectivity indirectly affects genetic differentiation through its effects on the frequency of the *Pgi-c* allele. Connectivity was included in the model as a latent variable – i.e. a construct that is not easily measured directly but can be indicated by a number of observed, and potentially correlated, variables that have some level of measurement error (Grace *et al.* 2012). We included connectivity as a latent variable, with the five connectivity hypotheses Si_metapop_, Si_water_, Si_forest_, Si_agriculture_, and Si_roads_ as indicators. We estimated model parameters using maximum likelihood and assessed model fit with chi-square tests, where a p-value *greater* than 0.05 indicates the model-implied covariance fits the observed covariance well (Grace *et al.* 2012). The relative effects of paths were evaluated from standardized path coefficients and individual significance of paths.

## Results

### Associations between landscape connectivity and Pgi

For data from 2011, selection on mixed effect models identified ten models with ΔAICc <2 (Table S4). All of these models included Si_forest_ and Si_water_, and seven of the models were embellishments of higher ranked models – i.e. they were the same as a higher ranked model but included uninformative parameters with little effect on model fit (Arnold 2010). We thus retained three models for further analysis and interpretation: the top model including an interaction of Si_forest_ and age and main effects of Si_water_ and Si_agriculture_, a model including an interaction of Si_forest_ and a main effect of Si_water_, and a third model including an interaction of Si_water_ and main effects of Si_forest_ and Si_agriculture_ (Table 1). All variables showed negative associations with *Pgi-c*, with Si_forest_ and Si_water_ having the strongest effects in new populations (Table 1). The effect of water and forest in new populations remained significant in latent factor mixed models as evidenced by a significant association of Si_water+forest_ with *Pgi-c* but no other loci (Fig. S2). In contrast Si_water_ showed a significant association with a putatively neutral locus but not *Pgi-c*, Si_forest_ showed a significant association with both *Pgi-c* and a different putatively neutral locus, and Si_water+forest+agriculture_ and Si_agriculture_ showed no significant associations with *Pgi-c* or any other locus (Appendix A; Fig. S2).

**Table 1.** Standardized regression coefficients showing the effects of connectivity and population age on the population frequency of the *Pgi-c* allele for the year 2011 and 2012. Candidate models with ΔAICc <2 and excluding uninformative parameters are shown (see Methods, and Table S4 and S5 for results of model selection). Variances explained by fixed effects (R^2^_m_) and jointly by fixed and random effects (R^2^_c_) are shown.

| Year | Standardized regression coefficients | | | | | | | | | $R^2_m$ | $R^2_c$ | $AIC_c$ | $\Delta AIC_c$ |
| --- | --- | --- | --- | --- | --- | --- | --- | --- | --- | --- | --- | --- | --- |
|  | (intercept) | age | Si<sub>water</sub> | Si<sub>forest</sub> | Si<sub>agriculture</sub> | Si<sub>roads</sub> | age:Si<sub>water</sub> | age:Si<sub>forest</sub> | age:Si<sub>roads</sub> |  |  |  |  |
| 2011 | 0.23 | -0.01 | -0.06 | -0.05 | -0.04 |  |  | -0.13 |  | 0.15 | 0.18 | -148.5 | 0 |
|  | 0.23 | -0.01 | -0.06 | -0.04 |  |  |  | -0.12 |  | 0.13 | 0.16 | -147.7 | 0.82 |
|  | 0.24 | -0.01 | -0.06 | -0.06 | -0.05 |  | -0.12 |  |  | 0.14 | 0.17 | -146.5 | 1.97 |
| 2012 | 0.24 | 0.006 | -0.05 |  |  | 0.02 | 0.08 |  | 0.11 | 0.10 | 0.10 | -216.4 | 0 |
|  | 0.24 | 0.003 | -0.05 |  |  | 0.02 |  |  | 0.13 | 0.08 | 0.09 | -215.2 | 1.17 |

The negative association between *Pgi* and connectivity in new populations switched in linear mixed models in 2012, and the effect of Si_forest_ disappeared (Fig. 2; Table 1). Model selection identified six models with ΔAICc <2 (Table S5). All of these models included an interaction of age and Si_roads_, and either a main or interactive effect of Si_water_. Four of the models were embellishments of higher ranked models. We thus retained two models for further analysis and interpretation: the top model containing interactions between of both Si_roads_ and Si_water_ with patch age, and a lower ranked model with an interaction between Si_roads_ and age and a main effect of Si_water_ (Table 1). Si_roads_ showed a strong positive association with *Pgi-c* in new populations, whereas Si_water_ showed a negative association with *Pgi-c* in old populations but only a weak or non-existent relationship in new populations in the top model (Table 1; Fig. S3). Neither Si_roads_ nor Si_water_ were found to have significant associations with *Pgi-c* or any other loci in latent factor mixed models (Appendix A; Fig. S2). Details on the calibration of the latent factor mixed models, including reports of genomic inflation factors can be found in Appendix A and Table S6.

**Figure 2.**
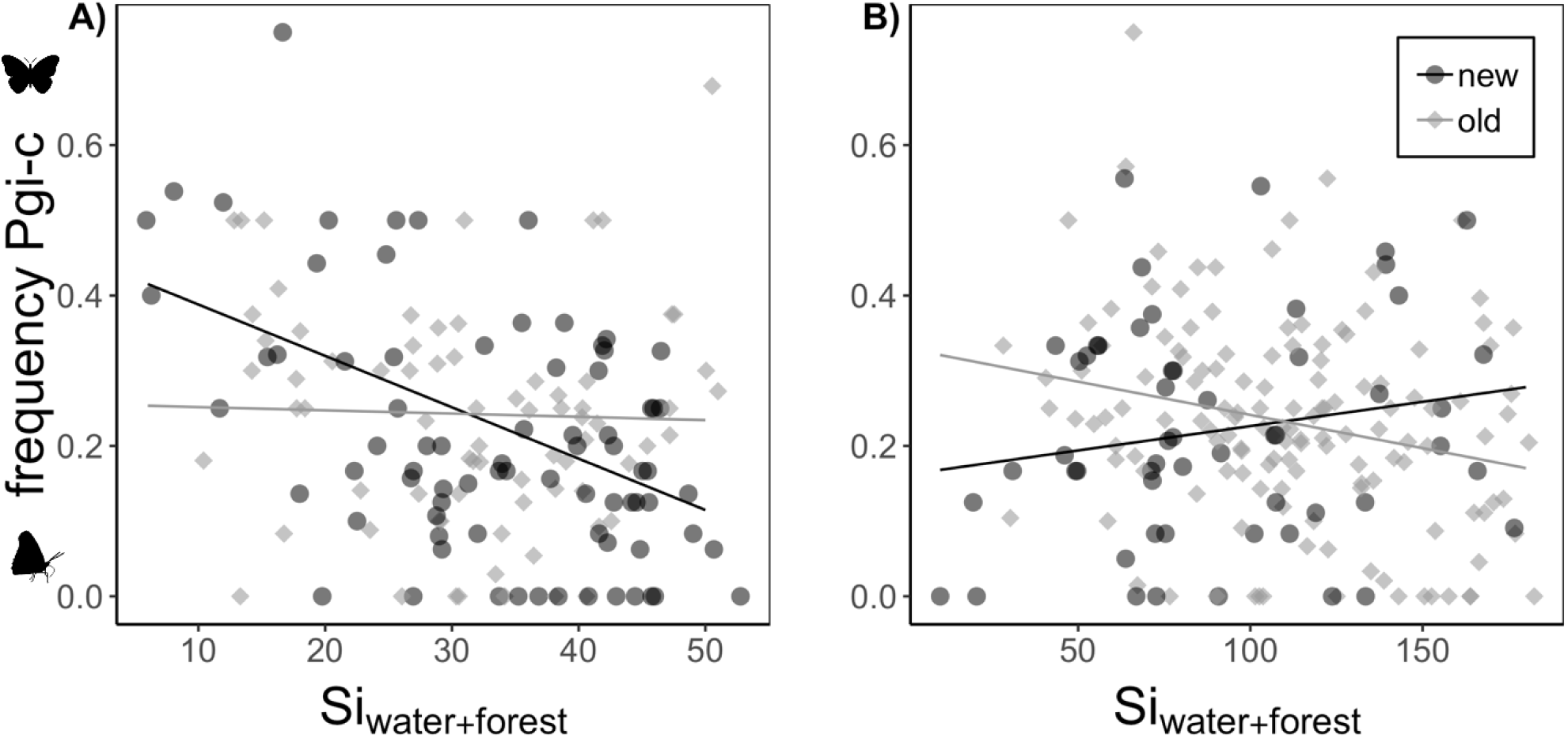
Scatterplots showing the relationship between the frequency of the *Pgi-c* allele and a patch connectivity metric incorporating forest and water as barriers to dispersal for the year 2011 when the metapopulation was recovering from a population decline (A) and 2012 when the metapopulation was at high density (B). Points represent local populations and old and new populations are shown in contrasting colours. Lines show predicted values of fixed effects from linear mixed models including Si_water+forest_ as a fixed predictor and genetic cluster as a random effect. See Fig. S5 for scatterplots showing variation across genetic clusters.

### Direct and indirect effects of landscape structure on genetic structure

For 2011, selection on mixed effect models testing for direct effect of connectivity and age on F_st_ identified a model containing only the random intercept as the most likely model (Table S7). We also tested the model after removing a single patch that had a very high F_st_ value. This led to the selection of a model including Si_water_ as the best, however it explained only 3% of the variation in F_st_, and the random intercept-alone model remained in the candidate set with ΔAICc <2 (Table S8). We thus conclude that there is no, or a very weak, direct effect of connectivity on genetic differentiation in 2011. For 2012, eight models had a ΔAICc <2, however five of them were embellishments of higher ranked models (Table S9). We thus retained three models: the top model including equal effects of Si_metapop_ and Si_water_, a model with just Si_metapop_, and a model with equal effects of Si_water_ and Si_roads_ (Table 2).

**Table 2.** Standardized regression coefficients showing the direct effects of connectivity on genetic differentiation (F_st_) for the year 2011 and 2012. Candidate models with ΔAICc <2 and excluding uninformative parameters are shown (see Methods, and Table S7 and S9 for results of model selection). Variances explained by fixed effects (R^2^_m_) and jointly by fixed and random effects (R^2^_c_) are shown.

| Standardized regression coefficients |  |  |  |  |  |  |  |  |
| --- | --- | --- | --- | --- | --- | --- | --- | --- |
| Year | (intercept) | $S_{i_{metapop}}$ | $S_{i_{water}}$ | $S_{i_{roads}}$ | $R^2_m$ | $R^2_c$ | $AIC_c$ | $\Delta AIC_c$ |
| 2011 | 0.08 |  |  |  |  | 0.02 | -464.5 | 0 |
| 2012 | 0.07 | -0.01 | -0.01 |  | 0.11 | 0.16 | -713.3 | 0 |
|  | 0.07 | -0.01 |  |  | 0.08 | 0.16 | -712.2 | 1.1 |
|  | 0.07 |  | -0.01 | -0.01 | 0.10 | 0.14 | -711.4 | 1.8 |

*Pgi* showed a significant association with connectivity only in new populations in 2011, and thus we only tested for indirect effects of connectivity on F_st_ (mediated by *Pgi*) in this partition. The full structural equation model including all five landscape connectivity indicator variables showed poor fit as indicated by a significant deviation of the observed covariance from the model-implied covariance (χ^2^=37.4, df=13, *p*<0.001). Because Si_metapop_ did not have strong effects in our earlier models, we removed it as an indicator variable. This model fit the data well as indicated by no significant deviation of the observed and modeled-implied covariance (χ_2_=11.94, df=8, *p*=0.15). The removal of Si_metapop_ did not affect the strength or significance of associations. Si_water_ and Si_forest_ were the strongest and only significant indicators of connectivity (Fig. 3). Our model supported a significant negative effect of connectivity on *Pgi-c*, and a significant negative effect of *Pgi-c* on genetic differentiation, but no significant direct effect of connectivity on genetic differentiation (Fig. 3; Fig. S4).

**Figure 3.**
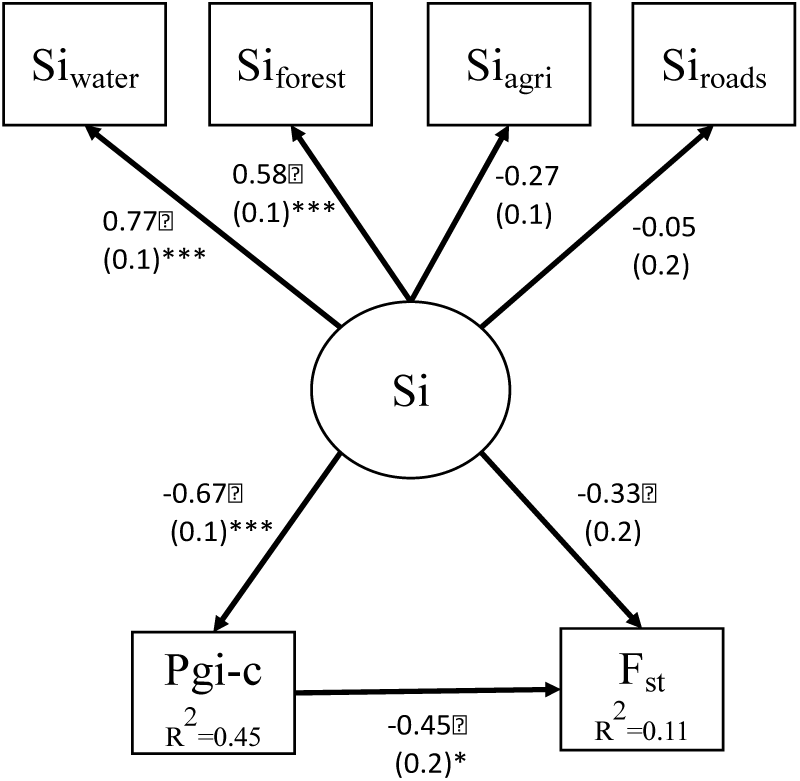
Results from a structural equation model test for direct and indirect effects of patch connectivity (S_i_) on genetic structure (F_st_) for new populations in 2011. Arrows show tested unidirectional relationship among variables. Connectivity (Si) was included as a latent variable described by four observed indicator variables, Si_water_, Si_forest_, Si_agriculture_, and Si_roads_. Standardized coefficients and associated standard errors are shown beside their respective paths, and significant paths are marked with asterisks. Variance explained is shown in the boxes of the endogenous variables, *Pgi-c* and F_st_.

## Discussion

Here we show that patch connectivity metrics incorporating landscape matrix best predicted the distribution of a dispersal polymorphism during a population expansion in the Glanville fritillary butterfly. In particular, newly colonized populations that were isolated by water and forest matrix had significantly higher frequencies of an allele associated with increased dispersal ability (*Pgi-c* allele). We further found that patch connectivity alone did not predict genetic differentiation at neutral markers, but rather the effect of landscape on genetic structure was mediated through individual variation in the *Pgi* locus; populations with higher frequencies of the *Pgi-c* allele had lower F_st_. In the following year when the density of populations increased, these relationships disappeared, suggesting that good dispersers only have an advantage when there are many empty patches to colonize. Together our results suggest that both individual variation in dispersal traits and landscape matrix heterogeneity are important for predicting spatial patterns of genetic variation.

### Landscape matrix predicts individual variation in dispersal

We found that spatial sorting of individuals based on their *Pgi* genotype was best explained by a connectivity metric that incorporated the effects of water and forest matrix in 2011. Importantly, the basic metapopulation model did not show a significant association with the frequency of *Pgi-c*. While previous work showed that metapopulation connectivity predicted the spatial sorting of individuals based on their *Pgi* genotype (Haag *et al.* 2005; Hanski and Saccheri 2006; Zheng *et al.* 2009), our results suggest that more complex processes are at play. There are a number of potential reasons for this. Previous patterns may have been confounded with demographic history. For example, we found evidence for a negative relationship between the frequency of *Pgi-c* and metapopulation connectivity for new populations, however, this relationship was *not* supported within individual genetic clusters (Fig. S5). Our sample size is also much larger and we were thus able to capture a larger amount of variation in landscape structure. In comparison, previous work selected population extremes (e.g. extremely low and high connectivity), and geographic distance might have been sufficient to capture patterns. It should also be noted that previous studies tested only a single model of patch connectivity, whereas we competed several models assuming different landscape structures.

Our results hence suggest that the *Pgi* dispersal polymorphism in the *M. cinxia* system in the Åland islands is not maintained by variation in patch configuration alone (i.e. the metapopulation model), but that the landscape matrix further influence dispersal. This is an important finding given that studies investigating the drivers of dispersal evolution almost exclusively use simplified landscape models that assume a homogenous matrix (Bowler and Benton 2005; Henriques-Silva *et al.* 2015). Knowledge of the importance of the landscape matrix is thus lacking, and this is one avenue in which landscape genetic approaches can contribute to understanding how the matrix might modify predictions of dispersal evolution. Our results suggest that landscape features that intervene discrete habitat patches matter, and this is unsurprising given that dispersal traits are often correlated with other aspects of species biology (Saastamoinen *et al.* 2018). For example, *Pgi* shows a genotype-by-temperature interaction in *M. cinxia*, where heterozygotes have higher flight metabolic rate (and thus dispersal ability) at moderate and cool temperatures, but individuals without a *c* allele fly better at very warm temperatures (Niitepold 2010). This might explain why forest is an important predictor of the spatial distribution of the dispersive allele, as it could provide a cooler environment for *Pgi-c* individuals. However, it is unclear if the association between forest and the frequency of *Pgi-c* is driven by individual differences in the use of the landscape matrix, or rather a distance effect – i.e. if *Pgi-c* individuals are able to fly further or faster through the forest.

### Individual variation in dispersal predicts genetic differentiation

In our test of direct effects of connectivity on genetic differentiation, we found no evidence that any of the connectivity metrics predicted F_st_ in 2011 - newly colonized patches with lower connectivity did not display higher genetic differentiation at neutral loci compared to highly connected (Table 2). Rather, patches with higher frequencies of *Pgi-c* had significantly lower F_st_ (Fig. 3; Fig. S4). Although *Pgi* explained only a small proportion of variation in F_st_, the effects of the landscape matrix on dispersal and gene flow would have been completely missed if individual variation at the *Pgi* locus was not included in our model. This highlights the importance for integration of intraspecific variation in dispersal traits in landscape genetic studies. While sex differences have recently been considered in landscape genetic models (e.g. Paquette *et al.* 2014; Bertrand *et al.* 2017), to our knowledge, only two papers have integrated non-sex related variation in dispersal traits: DiLeo *et al.* (2014) found that individual variation in the number of flowers on dogwood trees influenced spatial patterns of gene flow beyond the effects of Euclidean distance; and McDevitt *et al.* (2013) found that genetic admixture of weasels in Poland was likely mediated by the movement of medium-, rather than large-or small-bodied weasels. While we acknowledge that variation in dispersal traits are often hard to measure, the increasing accessibility of genomic data will facilitate the identification of candidate loci relevant to dispersal in non-model organisms (e.g. Swaegers *et al.* 2015; Dudaniec *et al.* 2018). Because dispersal is often controlled by multiple genes (Saastamoinen *et al.* 2018), applying landscape genomic methods that can capture the effects of polygenic adaptation will be important (e.g. redundancy and canonical correlation analysis; Rellstab *et al.* 2015 and reference therein). We further expect our approach to be relevant not only to systems where multiple dispersal strategies exist within a single landscape, but also the perhaps more common case of directional selection, where dispersal might be under positive or negative selection depending on broad patterns of landscape structure (Cheptou *et al.* 2017). An increasing number of studies have reported variability in landscape genetic relationships across replicate landscapes (e.g. Bull *et al.* 2011; Dudaniec *et al.* 2012; DiLeo *et al.* 2013; Balbi *et al.* 2018), and it will thus be interesting to see if divergent selection on dispersal genes might be a hidden source of variation contributing to these patterns (e.g. Peterson and Denno 1997).

Our study highlights the role of a single gene on the maintenance of gene flow across the landscape, and also joins a growing list of evidence that *Pgi* in particular has large effects on ecological processes in *M. cinxia* (reviewed in Niitepold & Saastamoinen 2018). However, it is unlikely that *Pgi* is acting alone. For example, work by Wheat *et al.* (2011) showed that *Pgi* may epistatically interact with other genes, such as succinate dehydrogenase (*Sdhd*); an allelic combination at these two loci was associated with maximal metabolic endurance in *M. cinxia*. Linkage of *Pgi* to other functional loci have not been resolved, but the low frequency and fitness of individuals homozygous for the C allele suggest possible linkage to a deleterious mutation (Orsini *et al.* 2009), and that the variation in Pgi is maintained through balancing selection via a heterozygote advantage (Wheat 2009). Further, females colonizing new populations have been found to be divergent in a suite of life-history traits, many but not all are associated with variation in *Pgi* (Hanski *et al.* 2006; Saastamoinen 2008; Kvist *et al.* 2013; Wheat et al. 2011). This suggest a more complex dispersal syndrome, the genomic architecture of which remains to be characterized. We further emphasize that *Pgi* explained only 10% of variation in F_st_, and clearly other processes are at play that may not have been captured by our model (e.g. temperature and condition-dependent dispersal).

### Context matters

Associations between patch connectivity and variation in *Pgi* in new populations in 2011 disappeared in 2012. Modeling studies on *Pgi* indicates that *Pgi-c* individuals should have the greatest selective advantage when there are many empty patches to colonize (Zheng *et al.* 2009; Hanski *et al.* 2011). Thus it was expected that our results would be much stronger in 2011 – a year that marked the largest expansion recorded in Åland following a large population decline that left many empty patches. In comparison, the metapopulation experienced a large increase in population size in 2012 but relatively fewer colonization events; all patches in 2012 had high connectivity. This appears to be driven by the much higher number of potential source patches and nests in sources in 2012, and less by difference in distances between sources and targets (Fig. S6). Observations from mark-recapture suggest that *M. cinxia* exhibit negative density dependent dispersal (Kuussaari *et al.* 1996), suggesting that there should be fewer dispersal events in the high density year 2012 compared to 2011. Intriguingly, effects of connectivity on *Pgi* even appeared to switch in 2012 (Table 1; Fig. S3), although these associations were not significant in latent factor mixed models. This might be suggestive of a more complex interaction between individual variation and density-dependent dispersal. Modeling work predicts that *Pgi-c* should rise to higher frequencies at very high population densities where it gains an advantage by spreading genes over more patches (Zheng *et al.* 2009), however this has not been empirically demonstrated. Future work should seek to resolve the drivers of yearly differences in dispersal, focusing particularly on the effects of density and weather, which may influence dispersive and non-dispersive genotypes differently.

Importantly, the association between *Pgi* and F_st_ in 2011 suggests that the polymorphism plays a key role in maintaining genetic variation across the landscape following perturbation. This finding provides a more mechanistic understanding of population persistence in this highly dynamic system. Recent work showed that regions in Åland with higher long-term frequencies of *Pgi-c* maintained higher metapopulation sizes, presumably by increasing colonization rates (Hanski *et al.* 2017). Our results suggest that regional persistence of the metapopulation might be further facilitated through *Pgi*-mediated genetic rescue.

### Model uncertainty

While some strong associations emerged from our analysis, model selection suffered from uncertainty with several likely, and sometimes non-nested models appearing to have similar support. This is a common problem with variables derived from landscape measures, which are inherently correlated (Smith *et al.* 2009; Prunier *et al.* 2015). Although our connectivity variables were well below typical collinearity thresholds (Dormann *et al.* 2013), it is likely that weak linear relationships still contributed to this uncertainty. This might also explain why connectivity variables with strong effects in one year did not emerge as important predictors in the next year, although part of this is also likely due to differences in spatial sampling of populations in the different years. While it is hard to say definitively which landscape features restrict dispersal, our results make a strong case for water as it was an important predictor across years, and forest as it had strong effects across multiple methods and different partitions of the data. What is less clear are the effects of other variables that were found to be important for prediction but of weak effect, with inconsistent results across methods (e.g. Si_agriculture_ in 2011). Future work would benefit from fine-tuning landscape resistance surfaces to better account for these potential small additive effects (e.g. using optimization; Peterman 2014), and from testing relationships under a broader set of conditions in carefully selected landscapes where the independent effects of landscape variables can be better teased apart.

### Conclusions

Our work adds to growing evidence that intraspecific variation plays a key role in driving diverse biological processes (Bolnick *et al.* 2011; Moran *et al.* 2016; Des Roches *et al.* 2018). We showed that heterogeneity in the landscape matrix is an important predictor of spatial variation in dispersal traits, and that this individual variation mediated the effects of landscape on genetic structure. Our results therefore highlight a need for better integration of studies on dispersal evolution and landscape genetics. While studies of dispersal evolution may need to consider more complex representations of landscape structure that captures heterogeneity in the landscape matrix, landscape geneticists should consider that key associations between landscape and genetic structure might be missed if intraspecific variation in dispersal is ignored.

## Acknowledgements

We thank Annukka Ruokolainen, Toshka Nyman, and Laura Häkkinen for sample preparation and DNA extraction and Swee Wong for post-processing of SNP data. We also thank Rolf Holderegger, Marie-Josée Fortin, Torsti Schulz, and Aapo Kahilainen for helpful conversations during the preparation of this manuscript. This study was funded by the Academy of Finland (Decision numbers 265641 to MS) and the European Research Council (Independent Starting grant no. 637412 ‘META-STRESS’ to MS).

## Author contributions

MFD analysed the data and wrote the first draft of the manuscript. AH and MS contributed substantially to the study design and revisions of the manuscript.

## Data accessibility

Data will be made available on DRYAD upon acceptance.

